# The mRNA-LNP platform’s lipid nanoparticle component used in preclinical vaccine studies is highly inflammatory

**DOI:** 10.1101/2021.03.04.430128

**Authors:** Sonia Ndeupen, Zhen Qin, Sonya Jacobsen, Henri Estanbouli, Aurélie Bouteau, Botond Z. Igyártó

**Affiliations:** Thomas Jefferson University, Department of Microbiology and Immunology, Philadelphia, PA

**Author notes:** Address correspondence to: Botond Z. Igyártó.

**Keywords:** mRNA-LNP vaccine, SARS-CoV-2 vaccine, LNP, inflammation, side-effects

## Abstract

Vaccines based on mRNA-containing lipid nanoparticles (LNPs) are a promising new platform used by two leading vaccines against coronavirus disease in 2019 (COVID-19). Clinical trials and ongoing vaccinations present with very high protection levels and varying degrees of side effects. However, the nature of the reported side effects remains poorly defined. Here we present evidence that LNPs used in many preclinical studies are highly inflammatory in mice. Intradermal injection of these LNPs led to rapid and robust inflammatory responses, characterized by massive neutrophil infiltration, activation of diverse inflammatory pathways, and production of various inflammatory cytokines and chemokines. The same dose of LNP delivered intranasally led to similar inflammatory responses in the lung and resulted in a high mortality rate.

In summary, here we show that the LNPs used for many preclinical studies are highly inflammatory. Thus, their potent adjuvant activity and reported superiority comparing to other adjuvants in supporting the induction of adaptive immune responses likely stem from their inflammatory nature. Furthermore, the preclinical LNPs are similar to the ones used for human vaccines, which could also explain the observed side effects in humans using this platform.

## INTRODUCTION

The nucleoside-modified mRNA-LNP vaccine platform used by Pfizer/BioNTech and Moderna in their SARS-CoV-2 vaccines has been widely tested in preclinical studies, and its effectiveness in supporting Tfh cells and protective humoral immune responses matches or surpasses other vaccines (Alameh et al., 2020). These vaccines’ mRNA component is nucleoside-modified to decrease potential innate immune recognition (Karikó et al., 2005, 2008). The LNP was chosen as a carrier vehicle to protect the mRNA from degradation and aid intracellular delivery and endosomal escape. The LNPs consist of a mixture of phospholipids, cholesterol, PEGylated lipids, and cationic or ionizable lipids. The phospholipids and cholesterol have structural and stabilizing roles, whereas the PEGylated lipids support prolonged circulation. The cationic/ionizable lipids are included to allow the complexing of the negatively charged mRNA molecules and enable the exit of the mRNA from the endosome to the cytosol for translation (Samaridou et al., 2020). Data support that some LNPs containing ionizable/cationic lipids are highly inflammatory and possibly cytotoxic (Samaridou et al., 2020). A preclinical study showed that mRNA complexed with Acuitas Therapeutics’ proprietary LNPs has adjuvant activity (Pardi et al., 2018a). However, the potential inflammatory nature of these LNPs was not assessed (Alameh et al., 2020; Pardi et al., 2018b, 2018a).

The human clinical trials of the Pfizer/BioNTech and Moderna vaccines have reported side effects often linked to inflammation, such as pain, swelling, fever, and sleepiness. (Jackson et al., 2020; Sahin et al., 2020; Walsh et al., 2020). Under the presumption that this vaccine platform is non-inflammatory, the reported side effects were interpreted as the vaccine being potent and generating an immune response. However, no studies have been undertaken to identify the potential causes of the local and systemic side effects. In this study, we took a systematic approach, focusing our attention on the injection site and analyzing the inflammatory properties of the LNPs used for preclinical vaccine studies (Awasthi et al., 2019; Laczkó et al., 2020; Lederer et al., 2020; Pardi et al., 2017a, 2017b, 2018c, 2018a). Using complementary techniques, we show that intradermal or intranasal delivery in mice of LNPs used in preclinical studies triggers inflammation characterized by leukocytic infiltration, activation of different inflammatory pathways and secretion of a diverse pool of inflammatory cytokines and chemokines. Thus, the inflammatory milieu induced by the LNPs could be partially responsible for reported side effects of mRNA-LNP-based SARS-CoV-2 vaccines in humans, and are possibly contributory to their reported high potency for eliciting protective immunity.

## RESULTS

### Intradermal inoculation with LNPs induces robust inflammation

mRNAs combined with LNPs were used in many preclinical studies and are key components of the recent Pfizer/BioNTech and Moderna SARS-CoV-2 vaccine (Alameh et al., 2020; Jackson et al., 2020; Sahin et al., 2020; Walsh et al., 2020). The mechanism of action of this mRNA-LNP platform is not well defined. The mRNA component is modified to decrease activation of interferon pathways (Karikó et al., 2005, 2008), but the mRNA complexed with LNPs was shown to have adjuvant activity (Pardi et al., 2018a). The mRNA-LNP platform promotes robust humoral immune responses, and humans receiving the vaccine often presented with typical side effects of inflammation, such as pain, swelling, and fever. (Jackson et al., 2020). Based on these observations, we hypothesized that mRNA-LNP adjuvant activity and the reported side effects in humans could stem from the LNPs’ inflammatory properties. mRNAs complexed with LNPs were used in preclinical studies at doses ranging from 3 to 30 μg/mouse (Laczkó et al., 2020; Pardi et al., 2018a). Therefore, we injected 10 μg (4 spots; 2.5 μg/spot) of these empty LNP formulated in phosphate buffered saline (PBS) or control PBS intradermally into adult wild type (WT) C57BL/6 (B6) mice. We sacrificed the mice at different time points post-injection and ∼1 cm^2^ skin samples from the injection sites were collected. The LNP-injected skin samples macroscopically showed signs of intense inflammation, such as redness and swelling (**Figure 1A**). Single-cell suspensions were prepared from these samples and analyzed for infiltrates using flow cytometry (**Figure 1B and Suppl. Figure 1**). Flow cytometry revealed massive and rapid leukocytic infiltrates dominated by neutrophils that slowly resolved by day 14 (**Figure 1B**). Removal of the ionizable lipid component from the LNPs abolished visible skin inflammation (**Figure 1C**) and the leukocytic infiltration (**Figure 1D**). Thus, LNPs used in preclinical studies promote swift inflammatory responses at the injection site, which depends on the ionizable lipid component.

**Figure 1.**
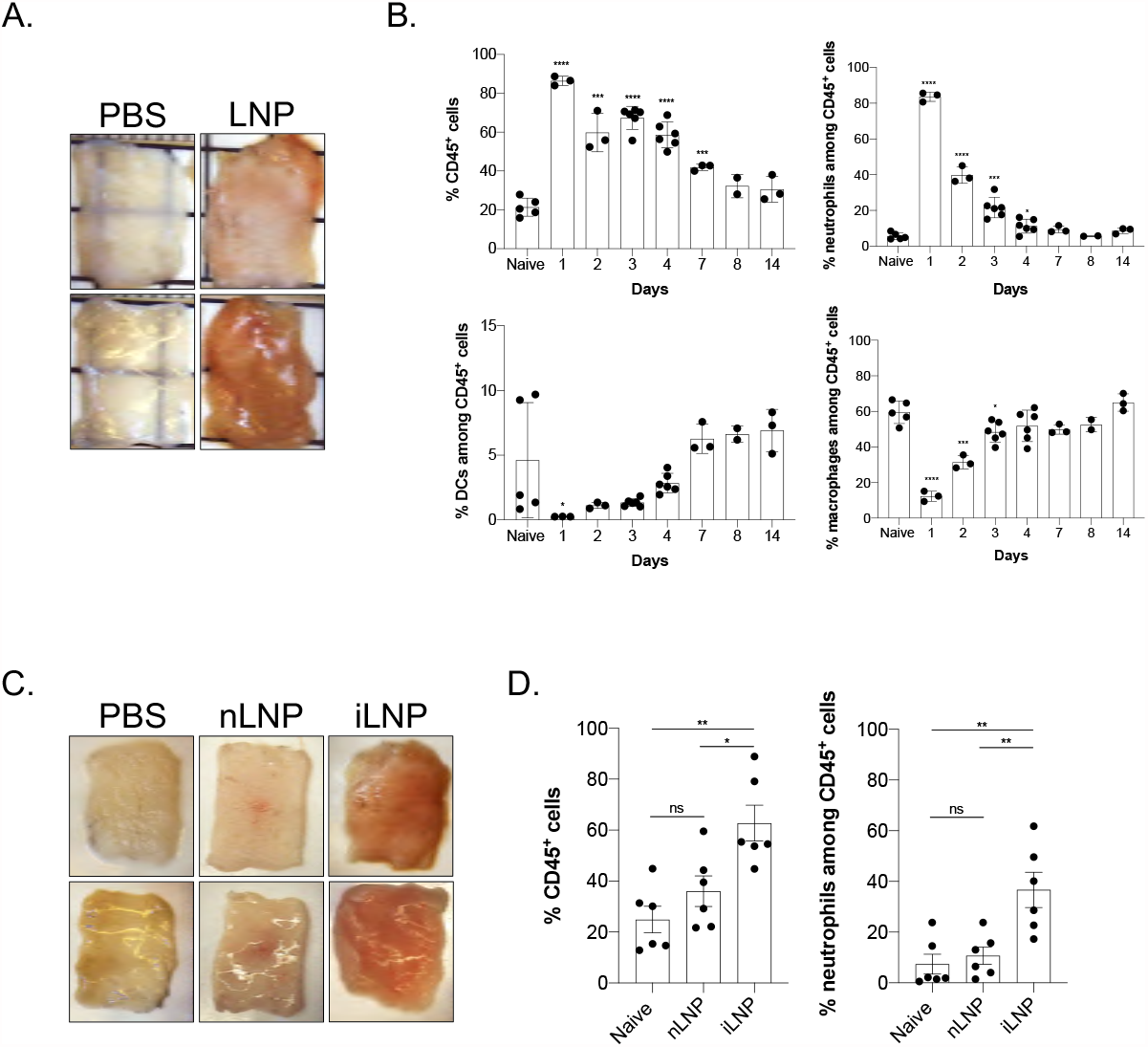
Intradermal inoculation with LNPs induces robust inflammation. **A**. Intradermal inoculation with LNP induced visible levels of inflammation. Pictures were taken 24 hours post PBS or LNP injection. **B**. Skin samples from the mice injected with PBS or LNP were harvested at the indicated time points, analyzed by flow cytometry, and displayed as cell percentages. **C**. As in **A**, but LNPs with (iLNP) or without (nLNP) ionizable lipids were used. Unlike iLNPs the nLNPs induced no visible signs of inflammation. **D**. Skin samples from **C** were analyzed for leukocytic infiltration 24 hours post inoculation. For all the charts the data were pooled from two separate experiments and displayed as percent ± SD. Each dot represents a separate animal. Student’s two-tailed t-test was used to determine the significance between naïve and the experimental samples. ****p<0.0001, ***p<0.0005, **p<0.005, *p<0.05, ns = not significant. No differences were observed between samples harvested from naïve or PBS treated animals and are used interchangeably throughout the manuscript.

To acquire a more in-depth understanding of the global changes triggered by the intradermal injection of the LNPs, we repeated the experiments presented above with LNPs complexed with control, non-coding poly-cytosine mRNA. Skin samples harvested one day after injection were split in two and analyzed using Luminex^®^ and bulk RNA-seq (**Figure 2A**). The Luminex^®^ data corroborated the flow cytometry findings and demonstrated the presence of a variety of inflammatory cytokines and chemokines (**Figure 2B, C and Suppl. Figure 2**), in comparison to control samples. Chemokines that attract neutrophils and monocytes and promote their functions, such as CCL2, CCL3, CCL4, CCL7, CCL12, CXCL1, and CXCL2, dominated the panel (**Figure 2B**). We further found large amounts of IL-1β, GM-CSF, IL-6, the signature cytokines of inflammatory responses (**Figure 2C**). Our RNA-seq analysis revealed that thousands of genes were upregulated (**Figure 2D**) upon LNP injection. With p<0.05 9,508 genes and with FDR<0.05 8,883 genes were differentially expressed. More importantly, confirming our flow cytometry and Luminex^®^ data, the genes associated with monocyte/granulocyte development, recruitment, and function (*Cxcl1, Cxcl2, Cxcl5, Cxcl10, Ccl2, Ccl3, Ccl4, Ccl7, Ccl12, Csf2, and Csf3*) and inflammation (*Il1b* and *Il6*) showed the highest fold increases over the control samples (**Figure 2E**). We also observed significant upregulation of gene transcripts associated with activation of inflammasomes, such as *Il1b* and *Nlrp3*, and downregulation of *Nlrp10*, which is known to inhibit inflammasomes (**Figure 2E**). Gene set enrichment analyses (GSEA) showed the activation of many different inflammatory pathways, including but not limited to viral infections, RIG-I, NOD-like, and Toll-like receptor signaling (**Figure 2F**). Pro-apoptotic and necroptotic gene sets were also significantly upregulated, as well as interferon signaling (**Figure 2F**).

**Figure 2.**
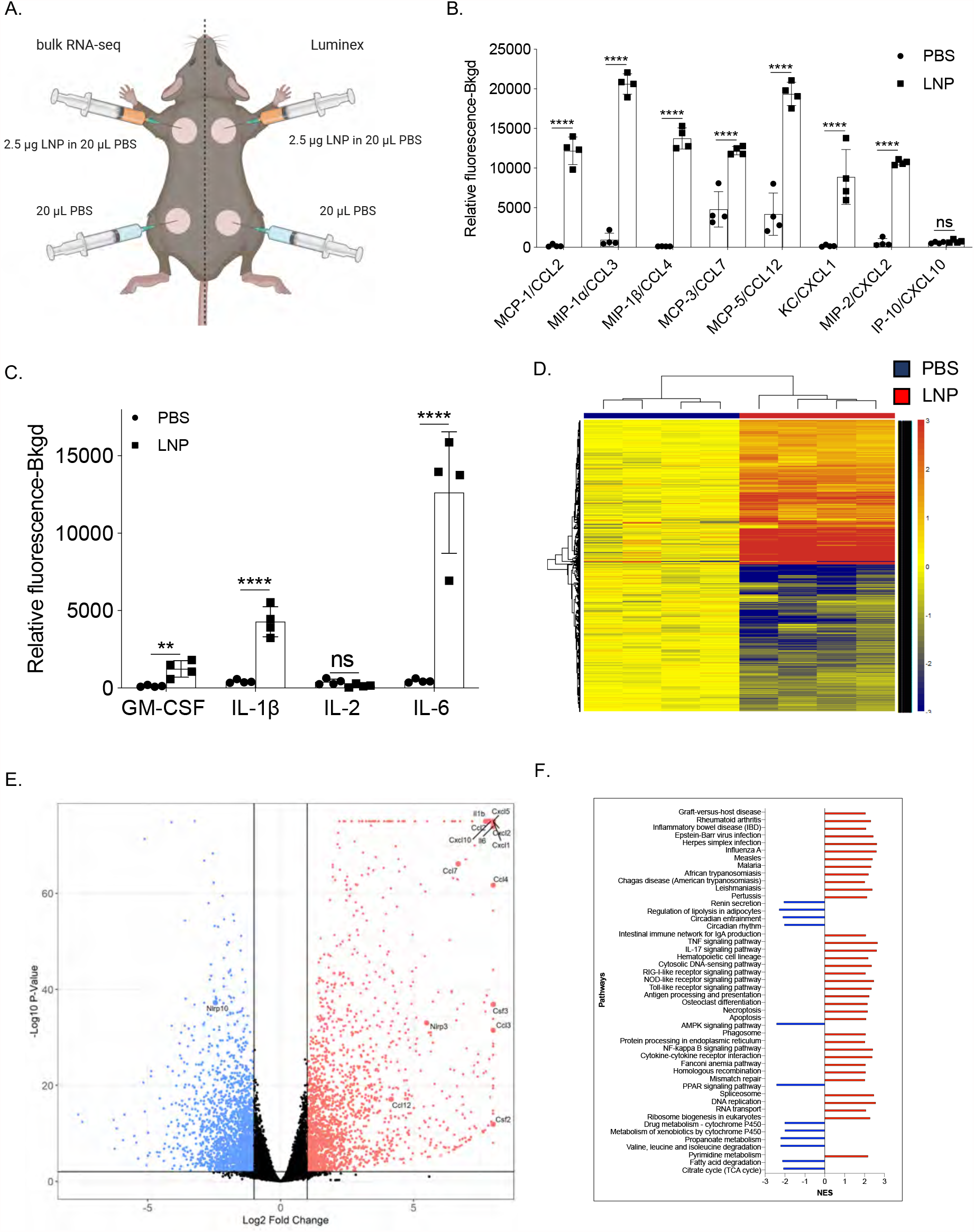
Intradermal inoculation with LNPs complexed with non-coding poly-cytosine mRNA leads to an inflammatory milieu. **A**. Experimental design. The mice were treated as indicated, and 24 hours later, the skin samples were prepared for Luminex^®^ and bulk RNA-seq analyses. **B. and C**. Luminex^®^ data summarizing inflammatory chemokines and cytokines induced by the LNPs. **D**. Heatmap of gene expression changes triggered by the LNPs (FDR < 0.05, log2 FC > 1 – 4091 genes). **E**. Volcano plot summarizing the up and downregulated genes upon LNP injection. **F**. GSEA analyses of the KEGG pathways and displayed as normalized enrichment score (NES). FDR<0.05. Pathways with NES less than ±2 are not displayed. N=4.

In summary, using different techniques, we show that LNPs, alone or complexed with control non-coding poly-cytosine mRNA, are highly inflammatory in mice, likely through the engagement and activation of various distinct and convergent inflammatory pathways.

### Intranasal inoculation with 10 μg of LNPs causes a high mortality rate in mice

To determine whether the LNP-induced inflammation is inoculation route independent, we further tested its potency in the airways. Mice are susceptible to intranasal inoculation of inflammatory compounds, and therefore it is a preferred route to test for side effects. We intranasally inoculated adult WT B6 mice with LNPs ranging from 2.5 μg to 10 μg/mouse and monitored their health and weight for up to eight days. We found that ∼80% of mice treated with 10 μg of LNP died in less than 24 hours (**Figure 3A**). The 5 μg dose killed ∼20% of the mice by that time, while all the 2.5 μg treated mice survived and showed no weight drop (**Figure 3B**) and no significant clinical signs of distress (**Figure 3C**). For the 5 and 10 μg doses, the surviving mice showed notable clinical scores of distress, such as shaking/shivering and they lost weight significantly during the first two days of treatment (**Figure 3B and C**). After the first ∼3 days, these mice did not continue to show significant clinical scores anymore, and their weight slowly started to normalize (**Figure 3C**).

**Figure 3.**
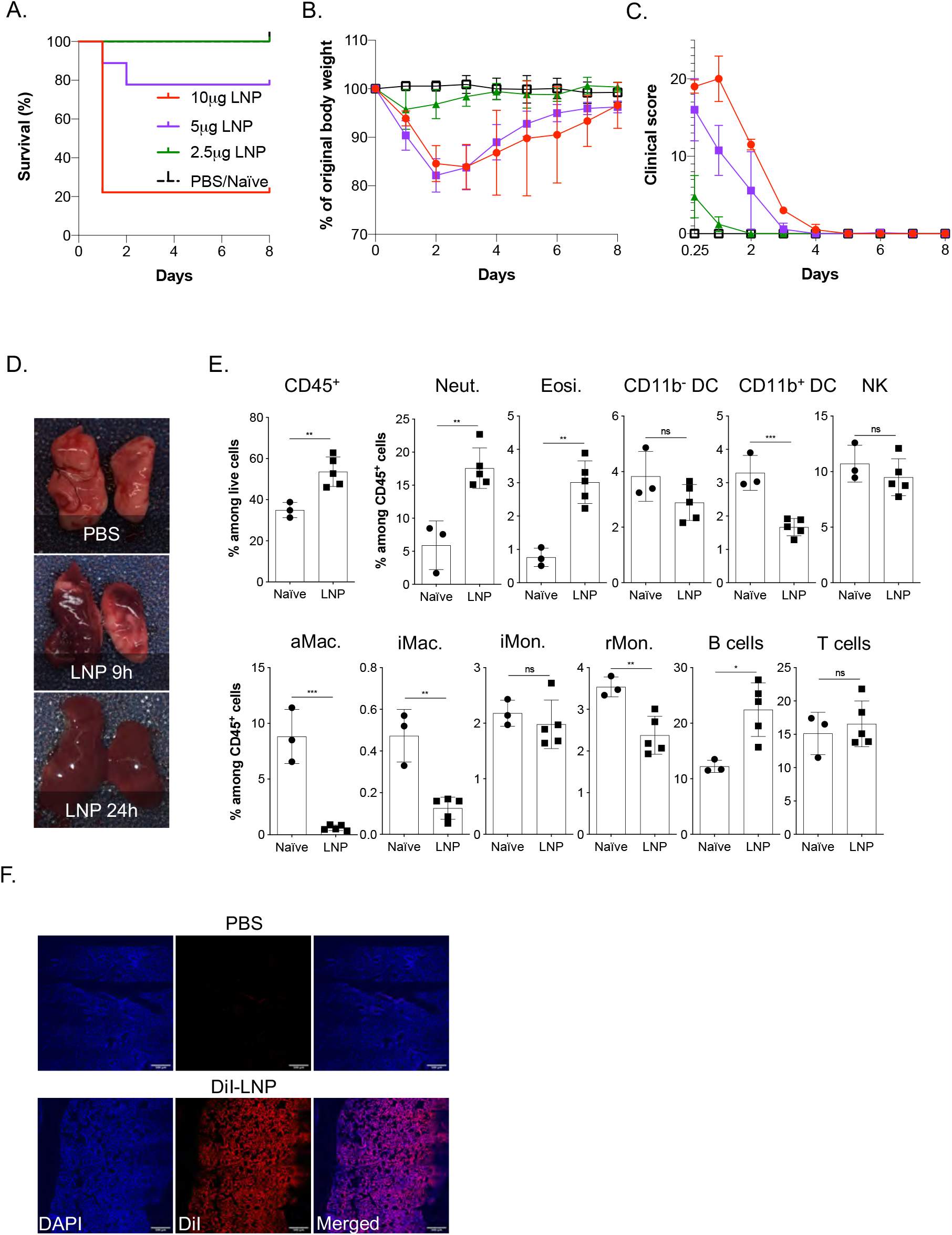
Intranasal LNP delivery induces robust lung inflammation and animal death. **A**. Animals were inoculated with the indicated doses of LNP and the survival rate, weight (**B**), and clinical scores (**C**) recorded daily for up to 8 days. Data was pooled from two independent experiments. N=9 for each group except PBS/Naïve where n=5. **D**. Lungs harvested at the indicated time points from PBS, and the 10 μg group were harvested and photographed. **E**. Animals injected with 10 μg of LNP were sacrificed 9 hours post inoculation and their lungs’ leukocytic composition determined by flow cytometry following a published gating strategy (Yu et al., 2016). Neut.= neutrophils, Eosi. = eosinophils, DC = dendritic cells, NK = natural killer, aMac. = alveolar macrophages, iMac. = interstitial macrophages, iMon. = inflammatory monocytes, rMon. = resident monocytes. **F**. LNP injection leads to fast and homogenous dispersion in the lung. Animals were inoculated with PBS or 10 μg of DiI-labeled LNP. Six hours later the lungs were harvested, prepared for histology, stained with DAPI and imaged using a confocal microscope. One representative image is shown. For all the charts the data were pooled from at least two separate experiments and displayed as percent ± SD. Each dot represents a separate animal. Student’s two-tailed t-test was used to determine the significance between naïve and the experimental samples. ***p<0.0005, **p<0.005, *p<0.05, ns = not significant.

We next tested whether intranasal inoculation leads to inflammation. For this, lung samples from PBS and 10 μg LNP treated mice were prepared for macroscopic analyses 9- and 24-hours post-inoculation and for flow cytometry 9 hours post-inoculation. In a matter of hours, the lungs turned red in color (**Figure 3D**). As was observed for the skin, flow cytometric analyses revealed significant leukocytic infiltration dominated by neutrophils and eosinophils, and a decrease in macrophages and certain DC subsets (**Figure 3E and Suppl. Figure 3**). To determine whether the LNPs reach the lung upon intranasal delivery, we inoculated mice intranasally with 10 μg of DiI-labeled LNPs. Six hours later, histology showed a homogenous distribution of the LNPs in the lung tissue (**Figure 3F**).

Thus, similar to skin inoculation, intranasal delivery of LNPs leads to massive inflammation. Furthermore, the LNPs’ inflammatory properties are not site-specific; and show a fast diffusion, dispersion and distribution rate in the tissues.

## DISCUSSION

Here we show that the LNPs used for many preclinical studies (Freyn et al., 2020; Laczkó et al., 2020; Lederer et al., 2020; Pardi et al., 2017b, 2018c, 2018a) are highly inflammatory. This could explain their potent adjuvant activity and their superiority, compared to other adjuvants, in supporting the induction of adaptive immune responses.

Previous preclinical mouse data suggested that mRNA complexed with LNPs have adjuvant activity (Pardi et al., 2018a). The mRNA is nucleoside-modified and purified specifically to decrease activation of certain innate inflammatory pathways (Karikó et al., 2005, 2008, 2011). Our injection-site-focused analyses revealed the inflammatory nature of these LNPs, which could provide a basis to their adjuvant properties. The cationic/ionizable lipid component of the LNPs are often inflammatory and cytotoxic (Samaridou et al., 2020). Indeed, we found that the proprietary ionizable lipid component of these LNPs is also inflammatory (**Figures 1C and D**). The mRNA-LNP vaccines are administered intramuscularly in humans. We reasoned that with the intramuscular delivery, the immediate inflammatory reactions in the deep tissue might remain hidden from the observer, which is especially true in mice covered by fur. Therefore, we opted to inject these constructs into the shaved skin intradermally or inoculate them intranasally. The rationale being that inflammation in the skin becomes visible to the naked eye, and substantial inflammation in the airways would trigger observable distress in the mice. Indeed, we found apparent signs of inflammation in both cases, supporting that the LNPs are inflammatory regardless of the route of inoculation. Thus, it is highly likely that the intramuscular injection of the LNPs triggers similar inflammatory responses in the muscle.

Humans present various side effects, most often pain, swelling, fever, and chills after intramuscular vaccination with the Pfizer/BioNTech or Moderna vaccines (Jackson et al., 2020; Sahin et al., 2020; Walsh et al., 2020). These are typical symptoms associated with inflammation triggered by cytokines such as IL-1β and IL-6 (Dinarello, 2018; Tanaka et al., 2014). Along with causing local inflammatory responses, these cytokines also act as major endogenous pyrogens (Conti, 2004) and instruct the hypothalamus to increase the body’s temperature (fever) to help overcome possible infections. In concordance with this, the intradermal inoculation of LNPs in mice led to the secretion of large amounts of major and minor pyrogens, IL-1β/IL-6 and macrophage inflammatory protein-α (CCL3) and macrophage inflammatory protein-β (CCL4), respectively (**Figure. 2B and C**). Furthermore, the observed activation of other inflammatory pathways and cell death could further accentuate the experienced side effects. However, further studies will be needed to determine the exact nature of the inflammatory responses triggered by mRNA-LNP vaccines in humans, and how much overlap there might be with the inflammatory signatures documented here for mice.

It remains to be determined how these LNPs or their ionizable lipid component activate distinct inflammatory pathways. In theory, LNPs could activate multiple pathways or alternatively engage only one that would initiate an inflammatory cascade. Some cationic/ionizable lipids bind and activate TLRs (Lonez et al., 2012, 2014; Samaridou et al., 2020; Tanaka et al., 2008; Verbeke et al., 2019). Our GSEA analyses revealed that these proprietary LNPs are also likely to activate TLR pathways, among others (**Figure 2F**). We also observed upregulation of inflammasome components such *Nlrp3* and enrichment of genes involved in necroptosis.

Inflammatory cell death, such as necroptosis and pyroptosis, could cause the release of DAMPs and the further enhancement of inflammation. Intranasal inoculation with LNPs led to a significant mortality rate, likely due to massive inflammatory responses induced in the lung (**Figure 3D and E**). However, some clinical symptoms, such as stiff tail, shaking, shivering, and lack of coordination (data not shown), also suggest brain involvement as a co-morbidity factor. LNPs as lipid particles can quickly diffuse (**Figure 3F**) and could potentially gain access to the CNS through the olfactory bulb or the blood. Whether these LNPs can cross the blood-brain barrier remains to be determined. However, they could still enter the CNS through the hypothalamus, which lacks the blood-brain barrier. Hypothetically, even LNPs injected into the periphery could reach the CNS through the blood, though the scant amounts reaching the CNS would likely not induce significant inflammation, but might still trigger hypothalamus-driven side effects such as fever, nausea and sleepiness.

People often present with more severe and systemic side effects after the booster shot. This raises the possibility that the adaptive immune response might somehow amplify side effects induced by the vaccine. One culprit identified so far is PEG, which is immunogenic. Antibodies formed against PEG have been reported to support a so-called anaphylactoid, complement activation-related pseudoallergy (CARPA) reaction (Kozma et al., 2020; Szebeni, 2005, 2014). Of note, since PEG is a compound frequently used in cosmetics and toothpaste, many individuals could have anti-PEG antibodies. We have discussed other possible mechanisms in our recent opinion article (Igyártó et al., 2021). Briefly, while mRNA mainly transfects cells near the injection site it could hypothetically reach any cell in the body (Maugeri et al., 2019; Pardi et al., 2015). The resulting translated protein could be presented on MHC-I in the form of peptides or displayed as a whole protein in the cell membrane. In both cases, cells with the vaccine peptide/protein on their surfaces could be targeted and killed by cells of the adaptive and innate immune system, CD8^+^ T and NK cells (via ADCC), respectively.

In summary, the first vaccination’s side effects, except for CARPA, are likely associated with robust inflammation induced by the LNPs. In contrast, after the second vaccination, side effects could be further exacerbated by immune responses targeting cells expressing the vaccine protein or its peptide derivatives. Whether innate memory responses (Netea et al., 2011) to LNPs also contribute to amplifying the side effects remains to be determined (**Figure 4**). Overall, the robust inflammatory milieu induced by LNPs, combined with presentation of the vaccine-derived peptides/protein outside of antigen-presenting cells, might cause tissue damage and exacerbate side effects. Since self-antigen presentation in an inflammatory environment has been linked to autoimmune disease development (Charles A Janeway et al., 2001), this merits further investigation, albeit not detected here.

**Figure 4.**
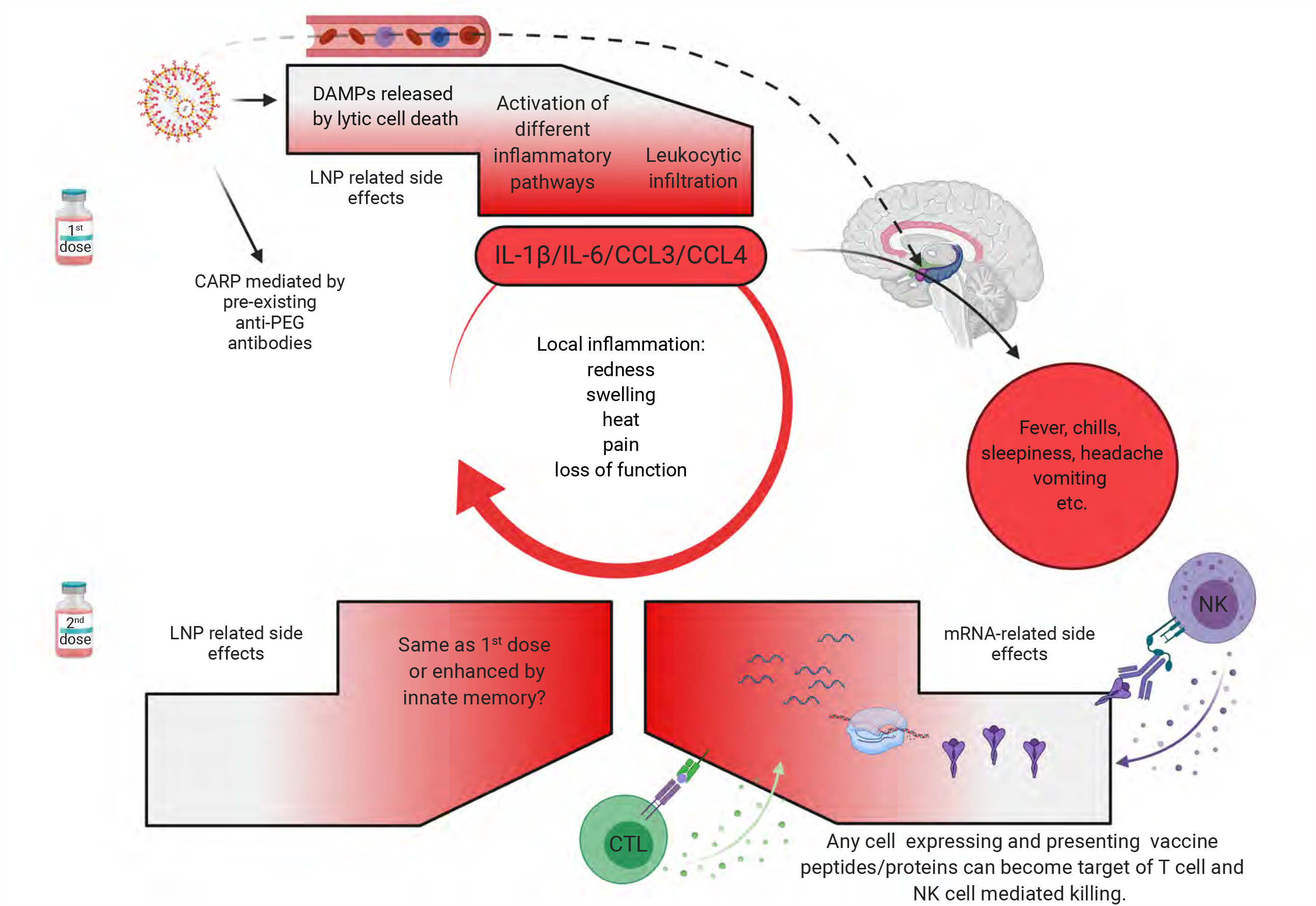
Potential mechanism of side effects. The side effects observed with the SARS-CoV-2 vaccine’s first dose are likely associated with the LNPs’ inflammatory properties. LNPs activate different inflammatory pathways that will lead to the production of inflammatory cytokines, such as IL-1β and IL-6 that can initiate and sustain local and systemic inflammations and side effects. LNPs might also diffuse from the periphery and reach any organs in the body, including CNS (hypothalamus) where they could directly induce side effects (dashed line). PEG is widely used as a food and medicine additive, and many of us develop antibodies to PEG. Therefore, the LNPs’ PEGylated lipids can induce CARP in humans with preexisting PEG-specific antibodies. Humans often experience more severe side effects with the second dose. Here we posit that might be due to multiple reasons. Firstly, innate immune memory against the LNPs might form after the first vaccination, and that could lead to even more robust inflammatory responses upon the second vaccination. Secondly, after the first vaccination adaptive immune responses are formed targeting the viral protein coded by the mRNA. As such, cells (shown as red shape) expressing the viral protein derived peptides or protein itself can become the target of CD8^+^ T or NK cell-mediated killing (ADCC), respectively. Since the LNPs could diffuse throughout the body and transfect any cell in their path with the mRNA, and the mRNA could also be further distributed through extracellular vesicles (Maugeri et al., 2019), the target population could potentially be vast and diverse.

mRNA-LNPs support very robust adaptive immune responses in animal models and humans (Alameh et al., 2020; Le Bert et al., 2020; Jackson et al., 2020; Sahin et al., 2020). The mechanism of induction of these immune responses so far has not been fully elucidated. Our findings revealed that LNPs used in some preclinical studies are inflammatory, which could explain their superiority to FDA-approved adjuvants in supporting the development of Tfh cells and humoral immune responses. Of note, their higher efficacy probably relies on activation of different inflammatory pathways exacerbated by direct cellular toxicity. The inflammatory properties of these LNPs’ should certainly be further exploited as an adjuvant platform in combination with proteins, subunit vaccines, or even in combination with existing attenuated vaccines (Bernasconi et al., 2021; Debin et al., 2002; Martins et al., 2007; Shirai et al., 2020; Swaminathan et al., 2016). LNPs, unlike other adjuvants, could thus serve a dual purpose, as both delivery vehicles for different cargos and as an adjuvant. However, it will be necessary to strike a balance between positive adjuvant and negative inflammatory properties as LNP-associated vaccines move forward. Since vaccine doses utilized in rodents are much higher than those used in humans(Nair and Jacob, 2016), detailed dose response studies will be required. Since some of DCs can support humoral immune responses in the absence of adjuvant, non-inflammatory LNPs might also be developed (Bouteau et al., 2019; Kato et al., 2020; Li et al., 2015; Yao et al., 2015).

## ACKNOWLEDGMENTS

This work was supported by departmental start-up funds and by R01AI146420 to B.Z.I. We thank the following core facilities for their help and assistance: genomic-, proteomics-, flow cytometry-, and the imaging core. Figures were generated using BioRender. The RNA-seq data can be found at the following GEO accession number: GSE167521.

## AUTHOR CONTRIBUTION

B.Z.I. conceptualized the study, interpreted the data and wrote the manuscript. S.N., and A.B. performed the intradermal mouse experiments and analyzed data. S.J. prepared figures. Z.Q. did the intranasal experiments and analyzed the resulting data. H.E. assisted with the microscopy.

## CONFLICT OF INTEREST

Authors declare no conflict of any sort.

## FIGURE LEGENDS

**Suppl. Figure 1**. Gating strategy for skin infiltrates data presented on Figure 1B.

**Suppl. Figure 2**. Extended cytokine and chemokine panel.

**Suppl. Figure 3**. (**A**) Gating strategy for lung leukocytes data presented on Figure 3E. (**B**). H&E stain of lungs harvested from PBS and 10 μg LNP treated mice 9 hours post inoculation.

## MATERIALS AND METHODS

### Mice

WT C57BL/6J mice of different ages and gender were purchased from Jax^®^ or bred in house. All experiments were performed with 6-12 weeks old mice. Mice were housed in microisolator cages and fed autoclaved food. Institutional Care and Use Committee approved all mouse protocols.

### Reagents

For our studies, we used an LNP formulation proprietary to Acuitas Therapeutics described in US patent US10,221,127. These LNPs were previously carefully characterized and widely tested in preclinical vaccine studies in combination with nucleoside-modified mRNAs (Laczkó et al., 2020; Lederer et al., 2020; Pardi et al., 2017a, 2018c, 2018a). The following LNP formulations were used: empty LNPs with or without proprietary ionizable lipid, DiI-labeled LNPs, and LNPs complexed with non-coding poly-cytosine mRNA.

### Intradermal inoculation and flow cytometry analyses

The day before injections, the hair from the back skin of adult WT B6 mice was removed using an electric clipper, and then the site of injections wet-shaved using Personna razor blades. The next day the mice were injected intradermally with 2.5 μg/spot LNPs in PBS (4 spots, 10 μg total) or PBS. At different time points post-injection, the mice were sacrificed and ∼1cm^2^ skin around the injection site harvested. The skin samples were then chopped into small pieces using a curved scissor and exposed to collagenase/hyaluronidase digestion as previously described (Kashem and Kaplan, 2018). Single-cell suspensions were stained for the following markers: fixable viability dye (Thermo Fisher), MHC-II, CD11b, CD11c, CD45, CD64, F4/80, and Ly6G (All from BioLegend). The stained samples were run on LSRFortessa™ (BD Biosciences) and the resulting data analyzed with FlowJo 10.

### Luminex^®^

The day before injections, the hair from the back skin of adult WT B6 mice was removed using an electric clipper, and then the site of injections wet-shaved using Personna razor blades. The next day the mice were injected intradermally with 2.5 μg/spot LNPs complexed with non-coding poly-cytosine mRNA or PBS (**Figure 2A**). Twenty-four hours later, the skin samples were collected and processed for Luminex^®^ and RNA-seq. The samples for Luminex were weighed and homogenized using a Dounce tissue grinder in 1.5 ml of 10 mM Tris pH 7.4, 150 mM NaCl, 1% Triton-X-100 per gram of tissue in the presence of Roche protease inhibitor cocktail. After incubation on ice for 30 minutes, the samples were spun at 10K RPM for 10 minutes at 4°C. The supernatants were harvested and filtered with a 0.22 µm Eppendorf tube filter (MilliporeSigma). The supernatants were aliquoted and kept at -80°C until further use. The samples were tested using the Bio-Plex Pro™ Mouse Chemokine Panel 33-Plex as per manufacturer’s instruction.

### RNA preparation, sequencing, data analyses and visualization

Total RNA was isolated from tissue lysates using the RNeasy Mini Kit (Qiagen), including on-column DNase digestion. Total RNA was analyzed for quantity and quality using the RNA 6000 Pico Kit (Agilent). Sequencing, data analyses and visualization were performed as we and others previously described (Kanehisa, 2000; Liberzon et al., 2011; Love et al., 2014; Su et al., 2020; Subramanian et al., 2005).

### Intranasal inoculation and flow cytometry analyses

Mice were anesthetized by intraperitoneal injection with a mixture of Xylazine/Ketamine. The LNPs were given intranasally at the doses of 2.5, 5 and 10 μg in 30 μL sterile PBS by placing droplets gently on the left nostril and allowed the mice to inhale. The clinical performances of the mice were scored daily for 8 days as described previously (Shrum et al., 2014). Besides, the body weight was also measured daily. Some of the mice from the 10 μg LNP dose and corresponding PBS controls were sacrificed at indicated time points post-inoculation, and lung samples were harvested for histology and flow cytometry. For histology, the samples were fixed in 4% PFA overnight and then embedded in OCT. Eight micrometer thick sections were prepared using a cryostat and counterstained with DAPI. Stitched confocal pictures were taken using a Nikon A1 microscope. The lung samples for flow cytometry were digested using the collagenase/hyaluronidase technique also used for the skin samples (Kashem and Kaplan, 2018). The resulting single-cell suspensions were stained with the following markers: fixable viability dye (Thermo Fisher), MHC-II, CD11b, CD11c, CD24, CD45, CD64, Ly6G and Ly6C (All from BioLegend) (Yu et al., 2016).

### Statistical analyses

All data were analyzed with GraphPad Prism version 9.0.0. Statistical methods used to determine significance are listed under each figure.

## Notes

### Competing Interest Statement

The authors have declared no competing interest.

