## Supplementary figures and images for "The mRNA-LNP platform’s lipid nanoparticle component used in preclinical vaccine studies is highly inflammatory"

### Suppl. Figure 1.

Upstream gates: singlets, live, CD45$^+$

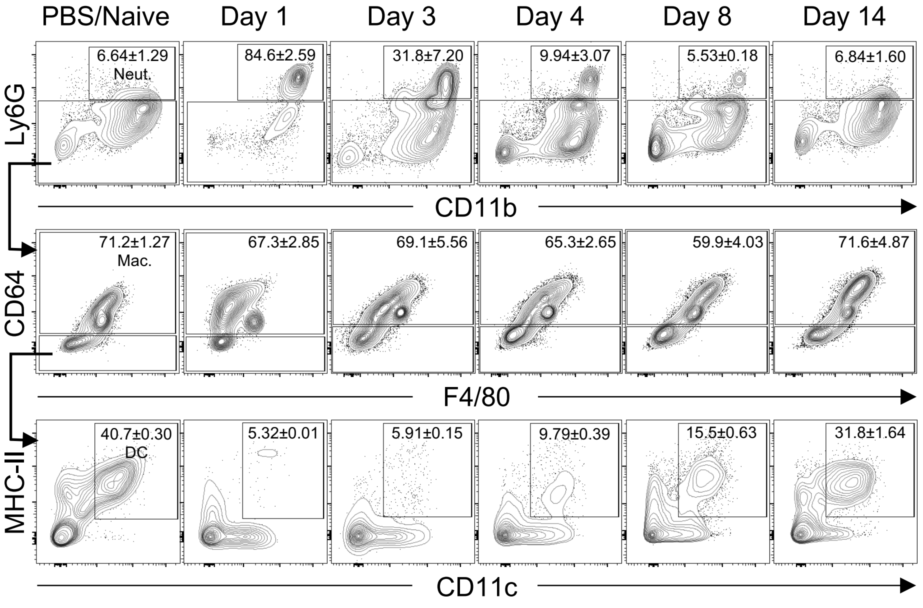

### Suppl. Figure 2.

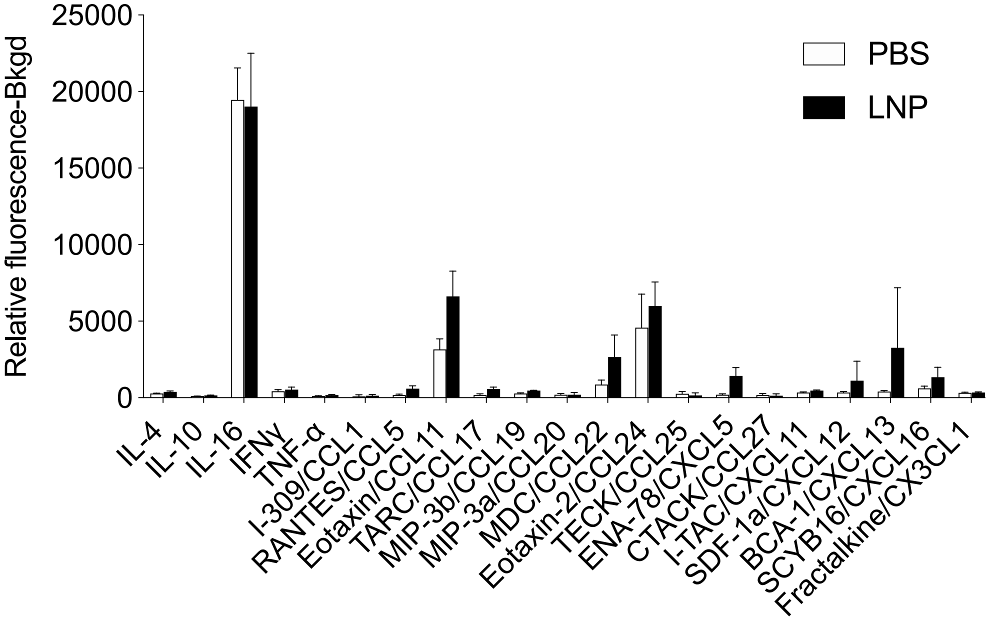

### Suppl. Figure 3.

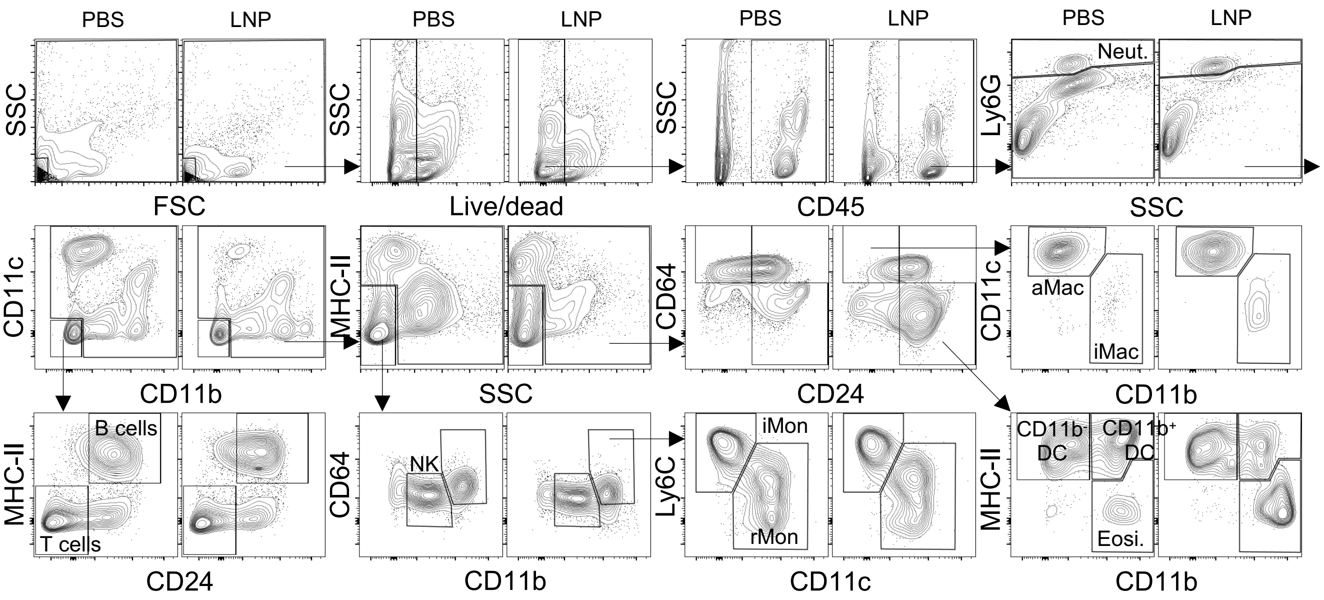

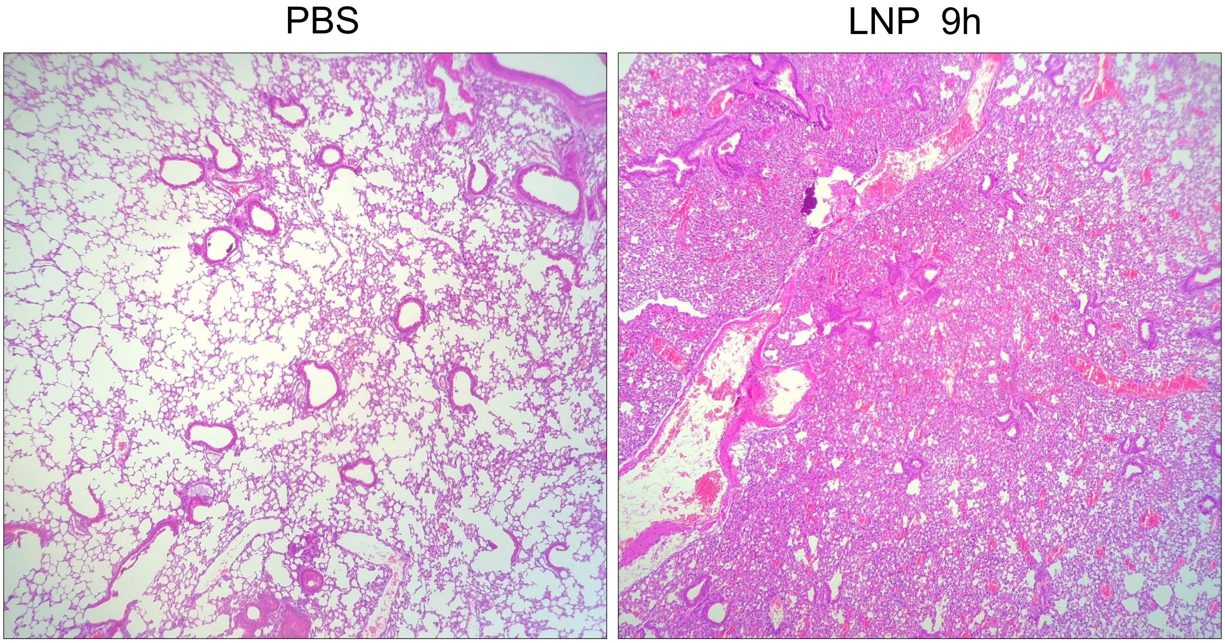
